# The AAA+ protein Msp1 recognizes substrates by a hydrophobic mismatch

**DOI:** 10.1101/2023.07.11.548587

**Authors:** Heidi L. Fresenius, Deepika Gaur, Baylee Smith, Brian Acquaviva, Matthew L. Wohlever

**Author notes:** Correspondence should be addressed to MLW.

## Abstract

An essential aspect of protein quality control is enzymatic removal of membrane proteins from the lipid bilayer. Failures in this essential cellular process are associated with neurodegenerative diseases and cancer. Msp1 is a AAA+ (ATPases Associated with diverse cellular Activities) protein that removes mistargeted proteins from the outer mitochondrial membrane (OMM). How Msp1 selectively recognizes and extracts substrates within the complex OMM ecosystem, and the role of the lipid bilayer on these processes is unknown. Here, we describe the development of fully defined, rapid, and quantitative extraction assay that retains physiological substrate selectivity. Using this new assay, we systematically modified both substrates and the lipid environment to demonstrate that Msp1 recognizes substrates by a hydrophobic mismatch between the substrate TMD and the lipid bilayer. We further demonstrate that the rate limiting step in Msp1 activity is extraction of the TMD from the lipid bilayer. Together, these results provide foundational insights into how the lipid bilayer influences AAA+ mediated membrane protein extraction.

## Introduction

A hallmark of eukaryotic cells is membrane bound organelles with each organelle characterized by unique protein and lipid compositions. As a foundational mechanism for organizing cellular reactions and signaling pathways, the cell devotes significant resources toward establishing and maintaining unique organelle proteomes and lipidomes. Despite the clear importance of targeting membrane proteins to the correct organelle, membrane protein trafficking and insertion pathways are surprisingly error prone and depend heavily on post-insertion quality control pathways to correct protein targeting errors.

One such pathway involves the AAA ATPase Msp1, which extracts proteins from the outer mitochondrial membrane (OMM) and peroxisome^1,2^. Msp1 substrates include endoplasmic reticulum (ER) tail anchored (TA) proteins that mislocalized to the OMM or peroxisomal TA proteins that are in stoichiometric excess of their binding partners^3^. Msp1 also helps relieve mitochondrial protein import stress by clearing stalled substrates from the TOM complex^4^.

ATAD1, the human homolog of Msp1, serves an expanded role in membrane proteostasis by regulating AMPA receptor internalization and selectively removing the pro-apoptotic protein BIM from the OMM^5,6^. Loss of Msp1/ATAD1 lead to failures in oxidative phosphorylation, impaired fear conditioning, apoptotic priming, and enhanced susceptibility to proteotoxic stress. Loss of ATAD1 in mice or humans is lethal^7,8^.

Given the clear health relevance, it is of paramount importance to understand how Msp1 recognizes substrates. In particular, it is unclear how Msp1 can selectively recognize mislocalized TA proteins from a large excess of properly localized mitochondrial proteins. There are currently several competing models in the literature for how Msp1 recognizes substrates^9,10^, however the role of the lipid bilayer in substrate recognition has been largely ignored. We previously used a reconstituted system to demonstrate that Msp1 is sufficient to recognize and extract a tail anchored protein from a fully defined lipid bilayer^11,12^. Furthermore, this reconstituted system showed physiological substrate selectivity as it specifically removed ER-TA proteins, but not the mitochondrial TA protein Fis1^6^.

Here, we use a new and improved extraction assay to test the role of the substrate TMD and lipid bilayer in substrate extraction. We vary TMD length and bilayer thickness to demonstrate that Msp1 senses a hydrophobic mismatch between the substrate TMD and lipid bilayer to select substrates for extraction. We also demonstrate that substrate extraction is modulated by changes in lipid bilayer fluidity, but not by increasing the thermodynamic stability of the cytosolic domain. This demonstrates that removal of the TMD from the lipid bilayer, rather than unfolding of cytosolic domain, is the rate limiting step in Msp1 activity. Together, these results demonstrate that the lipid bilayer plays in critical role in protein quality control and provide a detailed model for how Msp1 selectively recognizes and removes mislocalized membrane proteins from the far more abundant OMM resident proteins.

## Results

### Development of a more quantitative extraction assay

To observe potentially subtle differences in Msp1 activity, we completely redesigned the extraction assay. The new assay quantifies Msp1 extraction activity with the NanoLuc split luciferase system^13,14^. We cloned the 11-residue HiBiT sequence onto the C-terminus of our substrate, which is sequestered in the lumen of the liposome after reconstitution **(Figure 1A)**.

**Figure 1:**
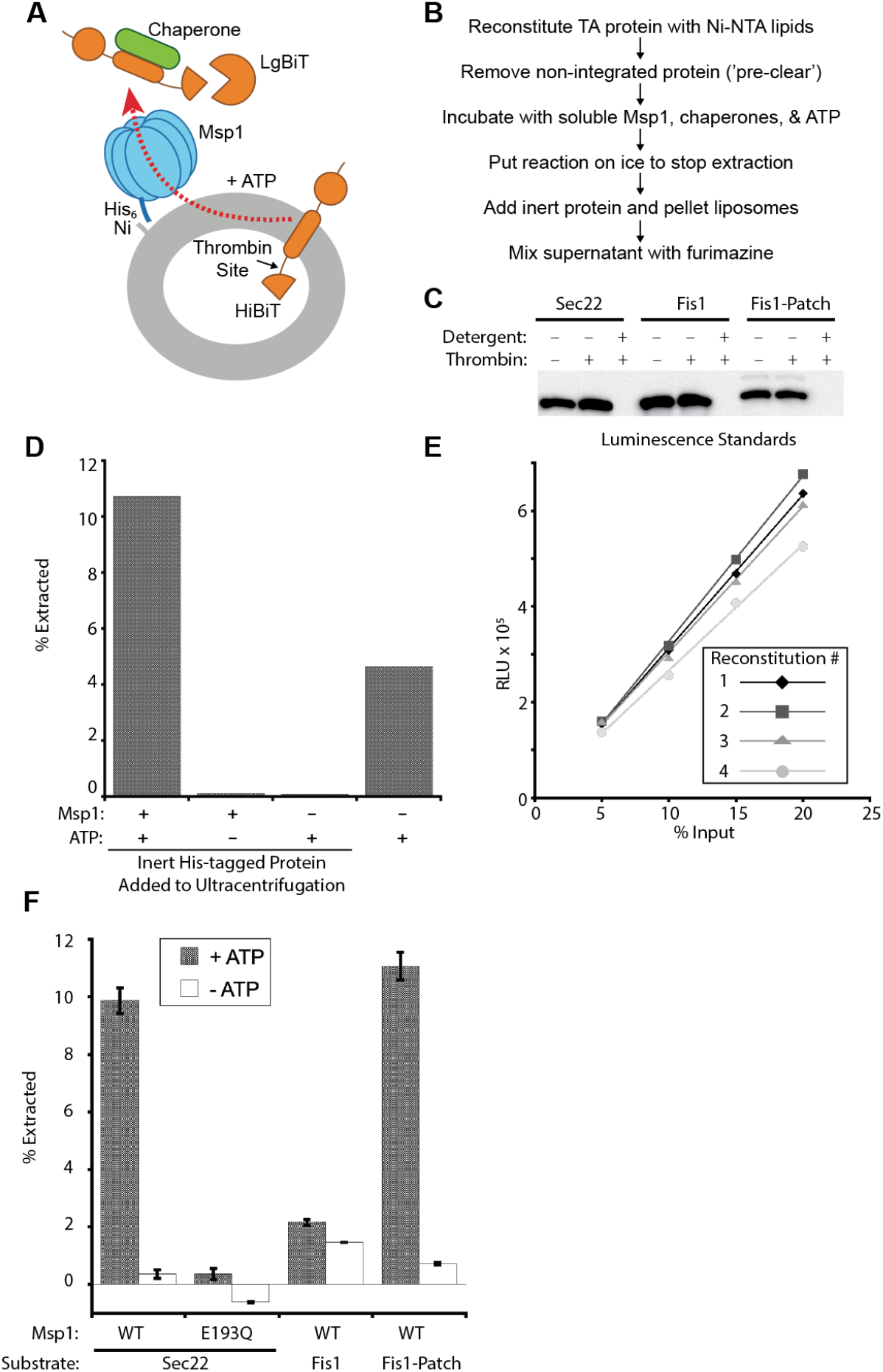
Validation of the split-luciferase based extraction assay. **A)** Diagram for split-luciferase extraction assay. Note that for properly reconstituted liposomes, both the HiBiT and thrombin sites are in the lumen of the proteoliposome. **B)** Workflow for split-luciferase assay. **C)** Protease protection assay shows that substrates are properly oriented. Anti-HiBiT western blot of pre-cleared liposomes shows minimal loss of signal upon addition of thrombin protease, but complete loss of signal with both thrombin and detergent. **D)** Addition of an inert His_6_-tagged protein to ultracentrifugation improves liposome pelleting efficiency and reduces background activity. **E)** Standard curves used to calculate percent substrate extracted. Four separate reconstitutions of the standard Sumo-Sec22 model substrate in standard liposomes show a linear and reproducible signal. Different concentrations of pre-cleared, but unreacted proteoliposomes were mixed with LgBiT and furimazine reagent in detergent, which permeablizes the proteoliposomes. The X-axis refers to the percent of substrate in the extraction assay. **F)** The split-luciferase extraction assay shows ATP-dependent and physiological substrate extraction activity. The known substrate Sec22 is extracted whereas the native mitochondrial protein Fis1 is not extracted unless a hydrophobic patch is added, which serves as an Msp1 recognition sequence. Error bars show standard error of the mean.

After reconstitution of substrate into pre-formed liposomes, the TA protein chaperones SGTA and calmodulin are added to remove any non-integrated substrate by immunoprecipitation **(Figure 1B)**. After this pre-clearing step, proper substrate orientation was validated by a protease protection assay that detects the HiBiT peptide **(Figure 1C)**. All of our substrates contain a thrombin protease site between the TMD and the HiBiT tag **(Figure 1A)**. Thrombin cleavage results in HiBiT peptide that is not resolvable by SDS PAGE/western blot. Addition of the thrombin protease led to a minimal loss of signal, whereas addition of both thrombin and the detergent Triton X-100 led to a complete loss of signal, demonstrating that the vast majority of substrate is properly oriented.

The assay is initiated by adding ATP. Fresh SGTA and calmodulin are also added to capture extracted substrates. After a 30 minute incubation at 30° C, Msp1 activity is inhibited by chilling the reaction on ice^12^ and then liposomes are pelleted by ultracentrifugation **(Figure 1B)**. We observed that the addition of an inert, His_6_-tagged protein (His_6_-MBP-ubiquitin) immediately prior to ultracentrifugation significantly reduces background signal, likely through enhanced liposome pelleting **(Figure 1D)**. The supernatant is then mixed with LgBiT and the luciferase substrate furimazine before luminescence is measured on a plate reader. To quantify total extraction activity, we generated a standard curve by mixing liposomes with LgBiT, the luciferase reagent furimazine, and detergent, which leads to membrane permeabilization **(Figure 1E)**.

Another major change is that we used soluble Msp1 instead of full-length Msp1. We and others had previously demonstrated that the Msp1 TMD can be completely changed with no effect on Msp1 activity *in vivo*^3,12^. Similarly, we also demonstrated that soluble Msp1 retains robust and physiologically selective extraction activity *in vitro* if it is anchored to liposomes via a Ni-His interaction^6^. The use of soluble Msp1 dramatically simplifies the reconstitution process as only the substrate protein needs to be reconstituted rather than co-reconstitution of substrate and six copies of Msp1. This allows us to directly compare extraction efficiency between different liposomes as we have eliminated any potential effects changes in lipid composition on Msp1 reconstitution efficiency.

We tested substrate extraction with our standard model substrate, Sumo-Sec22, and standard liposome preparation which mimics the outer mitochondrial membrane^15^. We observed robust, ATP dependent extraction **(Figure 1F)**. Extraction efficiency is ∼10% which is comparable to previously published results^6,11,12^. Compared to our previous western blot-based assay we observe significantly reduced signal in our negative controls. Importantly, the new assay design retains physiological substrate selectivity as the mitochondrial TA protein Fis1 is not extracted in our assay **(Figure 1F)**. However, addition of a known Msp1 recognition sequence, the hydrophobic patch from Pex15, leads to robust Msp1-dependent extraction consistent with previous results^6,16^. We conclude that the new assay has comparable extraction efficiency with our previously validated assay, but is faster, more quantitative, and has significantly improved signal to noise.

With a robust, quantitative, and physiological extraction assay now established, we next used this assay to examine the effects of the substrate TMD and lipid bilayer on Msp1 extraction activity. As proteoliposome reconstitutions are inherently variable, we took several steps to ensure experimental rigor and reproducibility. The extraction data involves at least 2 separate reconstitutions and between 2-4 technical replicates within each reconstitution. A new standard curve was generated for each new reconstitution. To ensure proper substrate reconstitution, a protease protection assay and a negative control (no ATP or ATPase deficient E193Q Msp1 mutant) was included in every assay.

### Msp1 extraction activity correlates with a hydrophobic mismatch between the substrate TMD and lipid bilayer

An outstanding question in the field is how Msp1 recognizes substrates. Our data clearly demonstrates that the hydrophobic patch is sufficient for substrate recognition. However, the juxtamembrane hydrophobic patch is unique to Pex15 and does not appear to be widely conserved across other Msp1 substrates. We have previously proposed that Msp1 recognizes substrates by a hydrophobic mismatch between the TMD of the mislocalized substrate and the lipid bilayer^9^.

To test this hypothesis, we first attempted to generate a hydrophobic mismatch by reconstituting the Sec22 model substrate into a series of liposomes with varying chain lengths. To minimize heterogeneity in the system, all liposomes were composed of phosphatidylcholine with both acyl chains containing a single cis double bond. The liposomes also contained 2% of DOGS-NiNTA, which was necessary to anchor soluble Msp1 to the liposomes. For simplicity, we will refer to the thickness liposomes as our T-series liposomes, with T1 referring to the thinnest liposomes (14:1, 14:1) and T5 the thickest liposomes (22:1, 22:1) **(Figure 2A)**.

**Figure 2:**
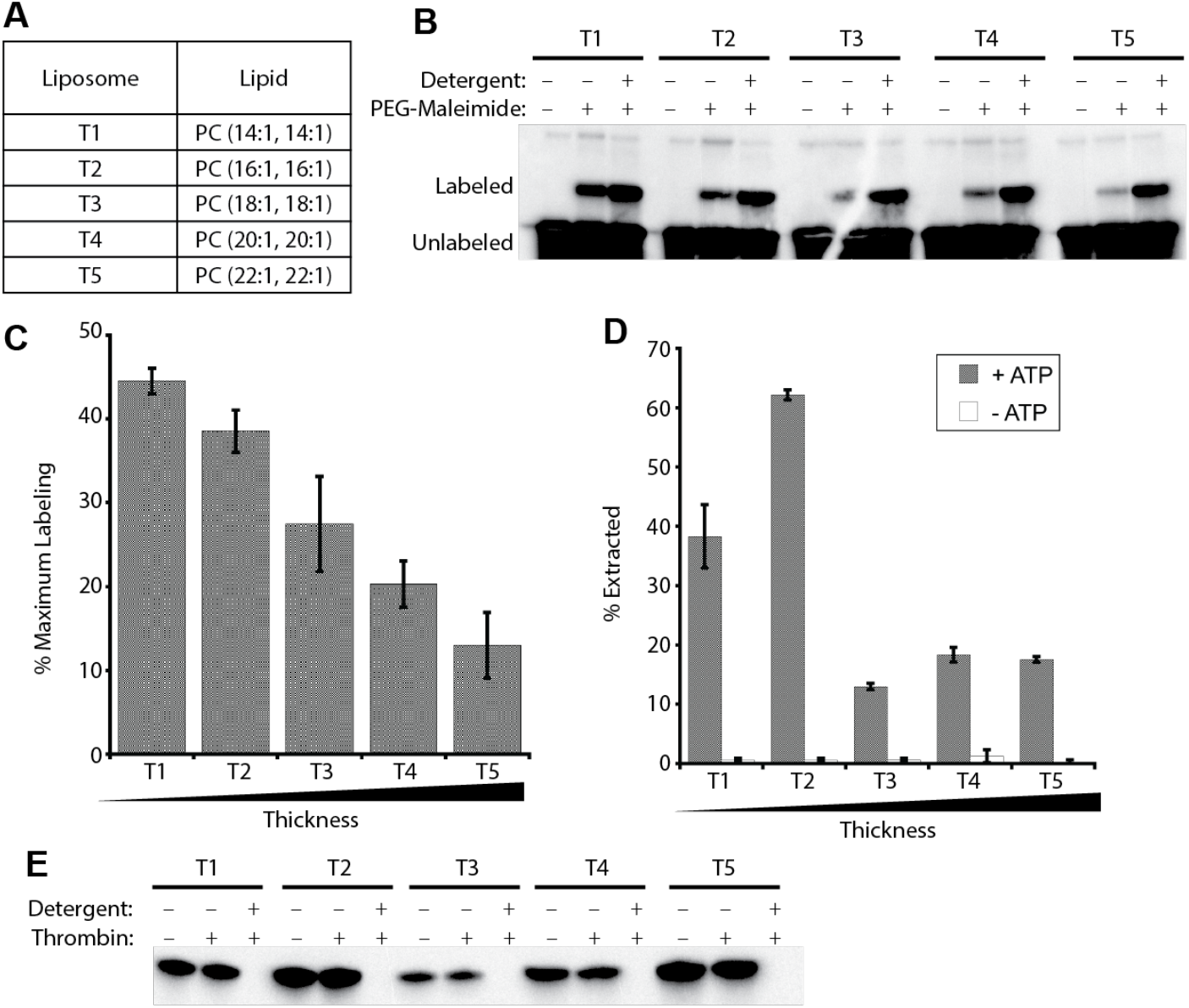
Msp1 extraction activity correlates with hydrophobic mismatch between the substrate TMD and lipid bilayer. **A)** Composition and thickness of T-series liposomes. **B)** PEG-maleimide labeling shows increased TMD exposure in thinner liposomes. **C)** Quantification of data from **B** from 3 separate reconstitutions. Error bars show standard error of the mean. The amount of labeled material in the + detergent sample is defined as 100% labeling. **D)** Extraction of the standard Sumo-Sec22 substrate in T-series liposomes. Msp1 shows more robust extraction activity in thinner liposomes. **E)** Protease protection assay of standard Sumo-Sec22 substrate in T-series liposomes.

We first sought to demonstrate increased exposure of TMD residues in a thinner lipid bilayer. To do this, we measured the solvent accessibility of a membrane-adjacent cysteine residue in our model substrate. We added a single cysteine residue immediately to the N-terminus of the Sec22 TMD. We then reconstituted the substrate into our T-series liposomes and reacted the liposomes with PEG_5000_-maleimide. If the cysteine residue is buried in the lipid bilayer, it will be protected from the membrane impermeable crosslinker. Conversely, a hydrophobic mismatch will lead to increased solvent exposure of the cysteine residue, resulting in increased labeling by PEG_5000_-maleimide, which can be easily observed by a size shift on SDS PAGE. As a positive control, the liposomes were solubilized with 1% Triton X-100 during labeling. We see a clear decrease in cysteine labeling as the model substrate moves to thicker liposomes **(Figure 2B & C)**.

Having demonstrated that thinner bilayers lead to increased solvent exposure of the TMD, we next asked if substrate extraction is enhanced in thinner bilayers. Consistent with our hypothesis, we observed significantly higher levels of substrate extraction in the thinner liposomes than the thicker liposomes **(Figure 2D)**. Unlike the linear drop in PEG-malemide labeling, the drop in substrate extraction appeared to follow a step function with robust extraction in the T1 and T2 liposomes and more modest extraction in T3-T5. We conclude that there is a strong positive correlation between hydrophobic mismatch of the substrate TMD with the lipid bilayer and Msp1 extraction activity.

### Substrates with a longer TMD show enhanced extraction activity

Many factors govern formation of a hydrophobic mismatch between a TMD and a lipid bilayer, including: TMD length and secondary structure, bilayer thickness, local membrane packing defects, and the tilt of a TMD relative to the bilayer plane^17–22^. In addition to enhanced hydrophobic mismatch in thinner bilayers, we hypothesized that the enhanced substrate extraction in the thinner bilayers could also be due to a lower thermodynamic barrier for substrate extraction^23^. Indeed, previous studies have shown that as the acyl chain length increases, the order of the lipid molecules increases^24^.

To determine whether the enhanced extraction in the thinner liposomes is due to increased hydrophobic mismatch or a lower thermodynamic barrier, we generated a hydrophobic mismatch by increasing the length of the TMD while holding the lipid bilayer composition constant. Increasing the length of the TMD should increase the stability of the substrate in the lipid bilayer, thereby increasing the thermodynamic barrier for substrate extraction^25,26^. If substrates with a longer TMD are extracted more robustly, it will be due to enhanced recognition by Msp1 rather than a lower thermodynamic barrier for extraction.

We therefore created a series of artificial substrates with TMD lengths ranging from 16-24 residues. For reference, Deep TMHMM predicts that the Sec22 model has a TMD length of 20 residues^27^. To prevent Msp1 interaction with specific sequences in the model substrate from potentially biasing our results, we designed artificial TMDs composed solely of leucine and alanine residues^28^ **(Figure 3A)**. To standardize the substrates, all TMDs were required to begin and end with leucine, have a Grand Average of Hydrophobicity (GRAVY) score of 3 +/-0.2, and contain 10-13 leucine residues^29^. We refer to these as our L-series substrates, with L16 having the shortest TMD at 16 residues and L24 having the longest TMD at 24 residues. The only difference between L-series substrates and the Sec22 model substrate is the TMD sequence and length.

**Figure 3:**
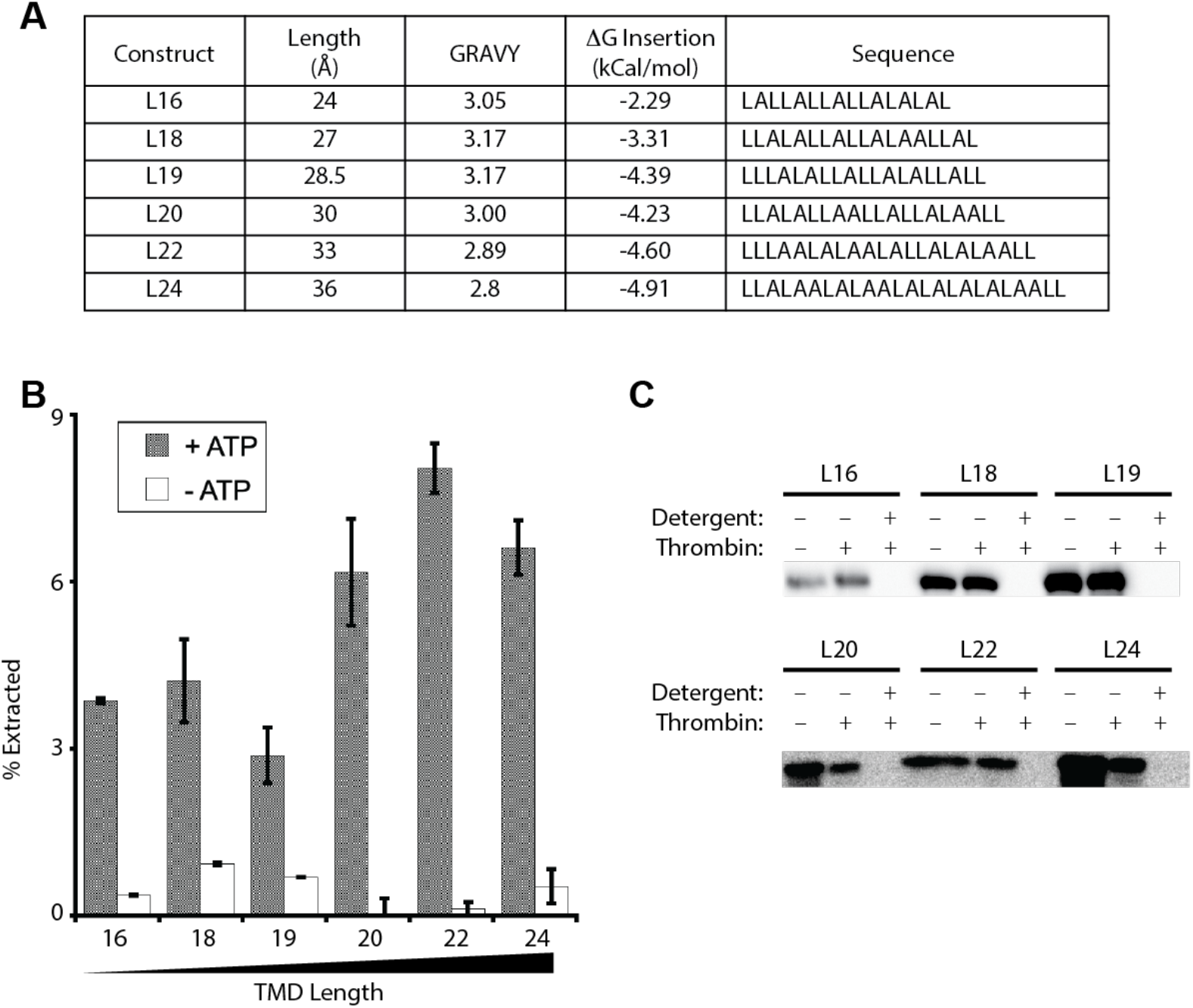
Design and extraction of L-Series Substrates. **A)** Design of L-series substrates. The length of the TMD is calculated by assuming a rise of 1.5 Å per residue. GRAVY score was calculated by inserting the TMD sequence into the GRAVY calculator (https://www.gravy-calculator.de/). The ΔG for TMD insertion was calculated using the website (https://dgpred.cbr.su.se/index.php?p=TMpred)^25,26^. TMD sequences for L-series substrates all start and end with Leucine. **B)** L-series substrates with a predicted hydrophobic mismatch show enhanced extraction activity. L-series substrates were reconstituted in standard liposomes. **C)** Protease protection assay of L-series substrates in standard liposomes.

We used an online calculator to predict the change in free energy for insertion of the L-series constructs into a model membrane^25,26^ **(Figure 3A)**. As expected, TMD insertion into the bilayer was more favorable with a longer construct. The thermodynamic barrier model predicts that the longer constructs should therefore have higher thermodynamic barriers for removal from the lipid bilayer and lower overall extraction. Conversely, the hydrophobic mismatch model predicts that the longer TMDs should lead to increased hydrophobic mismatch and therefore enhanced substrate extraction.

To distinguish between these two models, the L-series substrates were reconstituted into our standard liposomes and used in our extraction assay. All constructs showed ATP dependent extraction with the longer constructs showing higher levels of extraction than the shorter constructs **(Figure 3B)**. We again observed a step type function rather than a continuous change, with L20, L22, and L24 showing higher extraction than L16, L18, and L19 **(Figure 3B)**. Protease protection assays showed that the substrates were reconstituted in the proper orientation **(Figure 3C)**. While the change in extraction activity is relatively modest, it is significant and reproducible. The simplest explanation for the modestly enhanced extraction of the L20-L24 constructs is that a hydrophobic mismatch leads to enhanced substrate recognition, which is partially counteracted by increased thermodynamic stability of the substrate in the lipid bilayer.

### Variation of both substrate length and bilayer thickness supports hydrophobic mismatch model

Our standard liposomes contain a complex mixture of lipids which make it difficult to precisely predict the thickness of the hydrophobic core of the lipid bilayer. We also observed that a large discrepancy between TMD length and bilayer thickness led to low reconstitution efficiency. We hypothesized that these complications led to the non-linear trends in our extraction assays.

To test this hypothesis, we reconstituted the L-series substrates into the T-series liposomes, thereby gaining full experimental control over both TMD length and bilayer thickness.

We reconstituted the L20, L22, and L24 substrates into the T-series liposomes that most closely matched the predicted TMD length^24^. For the L20 substrate, this involved reconstitution into T1, T2, and T3 liposomes. Based on the length of the L20 TMD and the thickness of the T-series liposomes, there should be a hydrophobic mismatch in T1, a possible mismatch in T2, and no mismatch in T3 liposomes. Consistent with this model, we observed the most robust extraction in T1 proteoliposomes and the weakest extraction in the T3 liposomes **(Figure 4A)**. Similar trends were observed with L22 in T2-T4 liposomes **(Figure 4C)** and L24 in T3 and T4 liposomes **(Figure 4E)**. We conclude that a hydrophobic mismatch between a substrate TMD and the lipid bilayer is a major driver of Msp1 substrate selectivity.

**Figure 4:**
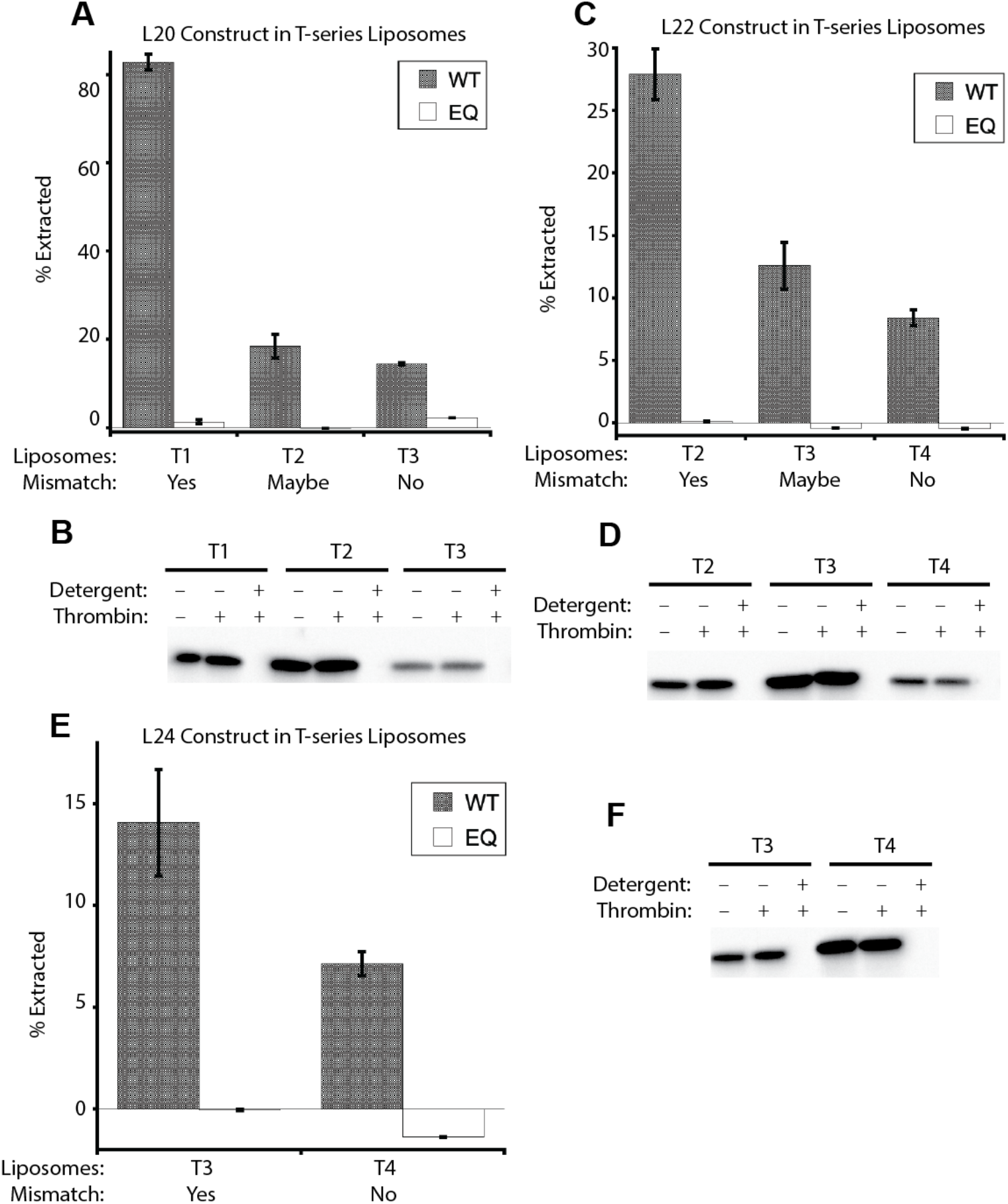
Extraction of L-series substrates in T-series liposomes. **A)** Extraction of L20 in is most robust when there is a predicted hydrophobic mismatch. L20 substrate was reconstituted in liposomes T1-T3. Based on the length of the TMD and thickness of hydrophobic core of the lipid bilayer^24^, there is a predicted hydrophobic mismatch T1, a possible mismatch in T2, and no predicted mismatch in T3. **B)** Protease protection assay for proteoliposomes in **A.** **C)** Similar to **A**, L22 substrate was reconstituted in liposomes T2-T4. There is a predicted hydrophobic mismatch T2, a possible mismatch in T3, and no predicted mismatch in T4. **D)** Protease protection assay for proteoliposomes in **C**. **E)** Similar to **A**, L24 substrate was reconstituted in liposomes T3 and T4. There is a predicted hydrophobic mismatch in T3 and no predicted mismatch in T4. **F)** Protease protection assay for proteoliposomes in **E**.

### Substrate extraction is more robust in bilayers with enhanced fluidity

Having firmly established that a hydrophobic mismatch between the substrate TMD and the lipid bilayer is the primary determinant of Msp1 extraction, we next sought to define the rate limiting step in substrate extraction. Previous studies with soluble Msp1 working on soluble substrates concluded that Msp1 is a processive unfoldase that recognizes long, unstructured tails at the end of a substrate^30^. This model predicts that Msp1 must first unfold cytosolic domains on a substrate before removing the TMD from the bilayer. In such a model, there are two major thermodynamic barriers that could serve as the rate limiting step in the reaction, 1) unfolding of the cytosolic domain or 2) extraction of the TMD from the lipid bilayer.

To test if TMD removal from the lipid bilayer is the rate limiting step in substrate extraction, we reconstituted our standard Sec22 model substrate into liposomes with increasing levels of unsaturated acyl chains. If unfolding of the cytosolic domain is the rate limiting step in substrate extraction, then increasing the fluidity of the lipid bilayer should have no effect on substrate extraction. Conversely, if the rate limiting step is removal of the TMD from the bilayer, we expect to see enhanced extraction activity in the more fluid membrane.

To minimize heterogeneity in the system, all liposomes were composed of phosphatidylcholine with a chain length of 18 **(Figure 5A)**. We will refer to the fluidity liposomes as our F-series liposomes, with F1 referring to the least fluid and F4 the most fluid. All liposomes were prepared in an identical manner and extruded through a 100 nm pore size. The molecular lipid packing densities were determined by using the fluorescent reporter C-Laurdan. The general polarization (GP) values followed the expected trend and were comparable to previously published values^31^ **(Figure 5B)**. Consistent with our hypothesis that TMD extraction is a rate limiting step, we observed a clear increase in substrate extraction with our most fluid liposomes **(Figure 5C)**.

**Figure 5:**
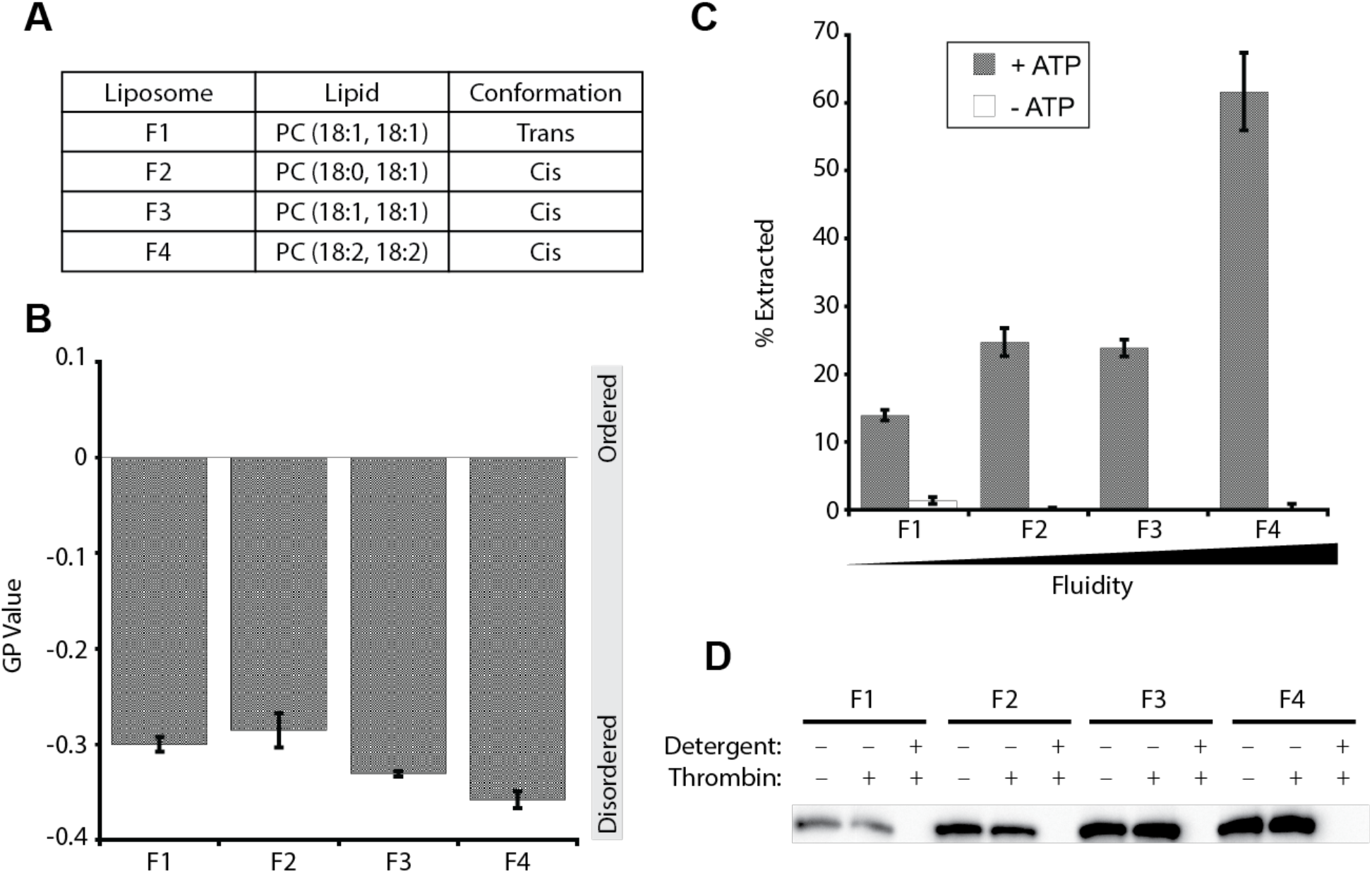
Msp1 extraction activity is enhanced with increasing membrane fluidity. **A)** Composition of fluidity series liposomes. **B)** Lipid packing of F-series liposomes measured by C-Laurdan spectroscopic measurements. Generalized polarization (GP) values range from +1 (most ordered) to -1 (most disordered). **C)** Extraction of standard Sumo-Sec22 substrate in F-series liposomes. **D)** Protease protection assay for proteoliposomes in **C**.

### Changing the stability of the cytosolic domain has no effect on substrate extraction

If TMD removal from the lipid bilayer is the rate limiting step in substrate extraction, then changing the thermodynamic stability of the cytosolic domain should have no effect on Msp1 extraction activity. To test this hypothesis, we replaced the Sumo domain in our model substrate with dihydrofolate reductase (DHFR) **(Figure 6A)**. Addition of the inhibitor methotrexate increases the resistance of DHFR to mechanical unfolding by AAA proteins^32^. We hypothesized that if TMD extraction is the rate limiting step in Msp1 activity, then addition of methotrexate should have no effect on substrate extraction.

**Figure 6:**
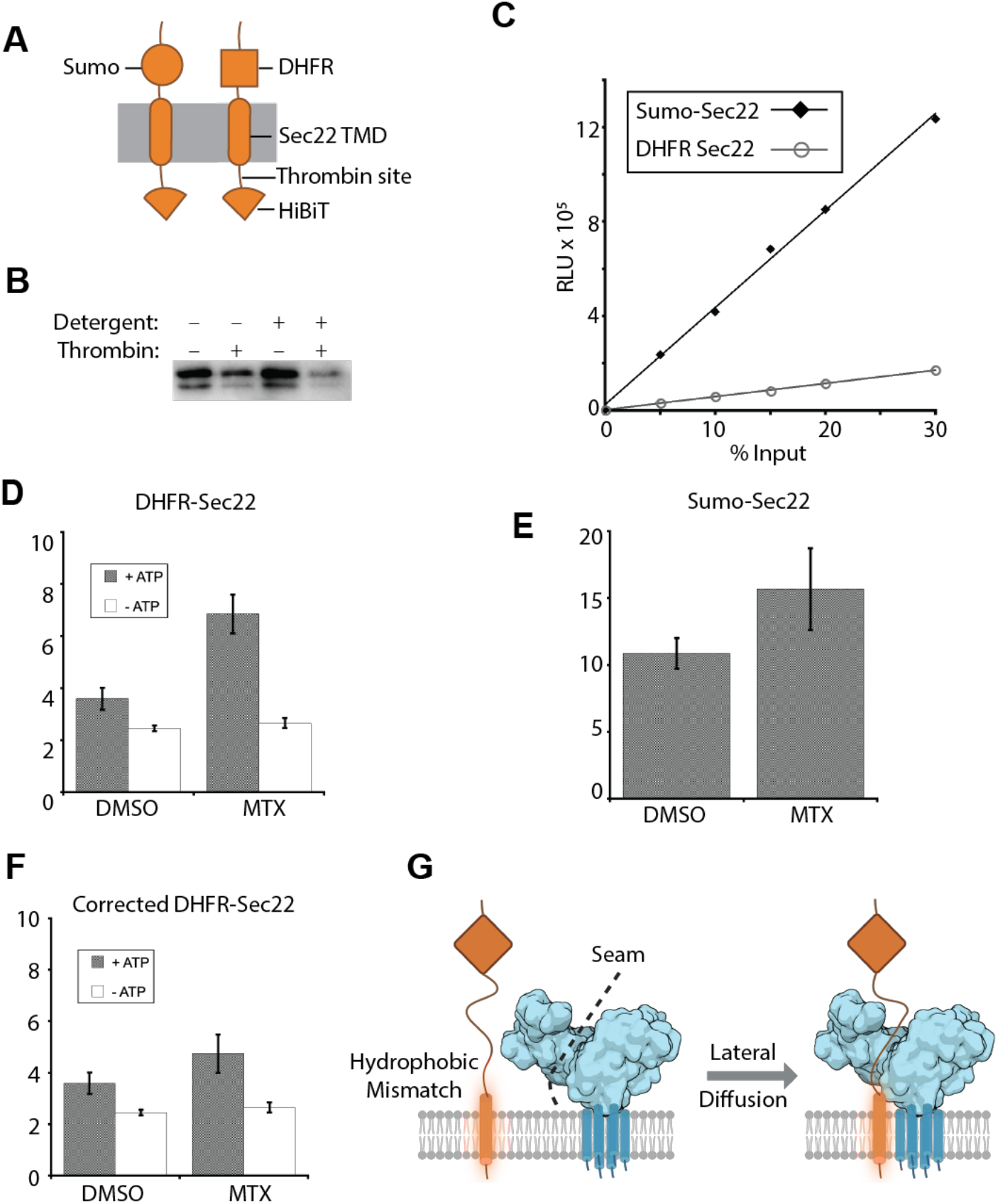
The rate limiting step in Msp1 activity is likely removal of the substrate from the lipid bilayer. **A)** Diagram of Sumo-Sec22 and DHFR-Sec22 model substrates. Note that the only difference is in the cytosolic domain. **B)** Protease protection assay for the DHFR-Sec22 substrate. **C)** Standard curves for reconstituted substrates show significantly lower levels of luminescence for DHFR-Sec22 than Sumo-Sec22 despite identical reconstitution conditions. This indicates that DHFR-Sec22 reconstitution is inefficient. Reconstitutions and standard curves were performed in parallel for both substrates. **D)** Extraction assay for DHFR-Sec22 model substrate shows ATP dependent extraction that is stimulated upon addition of methotrexate. **E)** Extraction assay for Sumo-Sec22 model substrate shows a 44% stimulation of extraction activity upon addition of methotrexate. **F)** Extraction of DHFR-Sec22 corrected for stimulating effects of methotrexate. Same data as in D, but extraction activity of the +ATP, +MTX sample was reduced by 44%. After this correction factor, the ATP dependent extraction is comparable to the DMSO negative control, indicating that methotrexate does not inhibit extraction of DHFR-Sec22. **G)** Diagram of the lateral diffusion model. Mislocalized ER-TA proteins have a hydrophobic mismatch with the OMM, leading to interaction between the solvent exposed TMD and hydrophobic residues in the seam of the Msp1 hexamer. The substrate enters the central pore of Msp1 by laterally diffusing through the seam, potentially bypassing the need for unfolding of cytosolic domains^45^.

We reconstituted the DHFR-Sec22 model substrate into our standard liposomes and performed the extraction assay in the presence of methotrexate or DMSO as a negative control. Overall extraction levels were lower for this substrate compared to Sumo-Sec22. As the sumo domain is a thermodynamically stable substrate with a higher T_M_ for thermal denaturation than DHFR^33,34^, we attribute this lower rate of extraction to technical challenges of reconstituting the poorly soluble DHFR-Sec22 substrate. Indeed, overall luminescence levels for DHFR-Sec22 substrate were only 10% of the Sumo-Sec22 substrate **(Figure 6C)** and the protease protection assay showed a higher proportion of substrate in the reverse orientation **(Figure 6B)**. Despite these technical challenges, we observed ATP dependent extraction of the DHFR-Sec22 substrate.

Surprisingly, addition of methotrexate increased the overall extraction of the model substrate compared to the DMSO only control **(Figure 6D)**. To test if methotrexate is enhancing Msp1 activity, we repeated the extraction assay with the Sumo-Sec22 model substrate, which should not interact with methotrexate. We observed a 44% increase in Msp1 activity in the presence of methotrexate **(Figure 6E)**. We then reanalyzed the extraction data by reducing the activity of the +ATP, +MTX sample by 44%. After this correction factor, we observed ATP dependent extraction of DHFR-Sec22 that is indistinguishable from the DMSO negative control **(Figure 6F)**, suggesting that methotrexate does not inhibit extraction of DHFR-Sec22. As changing the thermodynamic stability of the soluble domain has no effect on substrate extraction whereas increasing bilayer fluidity enhances substrate extraction, we conclude that the rate limiting step in Msp1 activity is extraction of the TMD from the lipid bilayer.

## Discussion

An essential aspect of membrane proteostasis is the removal of substrates from the membrane. However, our mechanistic understanding of this process, particularly the role of the lipid bilayer is incomplete. To address this knowledge gap, we developed a rapid and quantitative assay to measure Msp1-mediated substrate extraction from a lipid bilayer. We demonstrated that this assay has comparable extraction efficiency to our previously published assay, but with the added benefits of greater quantitation, faster turnaround time, and reduced background signal. Importantly, this system retains physiological substrate selectivity. Using this fully defined system, we systematically varied features of our model substrates and the lipid bilayer to understand the molecular basis for how Msp1 recognizes and removes substrates from the lipid bilayer. We demonstrated that Msp1 recognizes substrates through a hydrophobic mismatch between the substrate TMD and the lipid bilayer. Furthermore, we demonstrated that the rate limiting step in substrate extraction is removal of the TMD from the lipid bilayer rather than unfolding of the cytosolic domain.

Previous work had to led to the proposal of several different models for how Msp1 recognizes and extracts substrates. Weir et al. demonstrated that Msp1 extracts Pex15 from the peroxisome when it is in stoichiometric excess of its binding partner, Pex3^3^. This led to a model that orphaned substrates are less thermodynamically stable than a mature complex and/or expose an Msp1 recognition motif. However, subsequent studies demonstrated that Pex3 can directly inhibit Msp1 activity^30^. Using a soluble Msp1 construct that was artificially hexamerized by fusion to the PAN N-domain, Castanzo et al. demonstrated that Msp1 recognizes long, unstructured polypeptides^30^. However, the rate of substrate processing was quite slow, with a V_Max_ of 0.1 substrates hexamer^-1^ min^-1^. It should also be noted that these studies used soluble (ΔTMD) Msp1 and soluble substrates in the absence of membranes, so the physiological significance is unclear. Both of these models suggest that substrate thermodynamic stability plays a major role in regulating Msp1 activity.

Li et al. used photo-crosslinking to demonstrate that a juxtamembrane hydrophobic patch is critical for Msp1-Pex15 interactions^16^. Addition of this hydrophobic patch to the mitochondrial TA protein Fis1 is sufficient to convert it into an Msp1 substrate. This is a particularly appealing model as recent cryo-EM structures of *C. thermophilum* Msp1 and human ATAD1 show an exposed hydrophobic pocket within the N-domain that could provide a membrane adjacent binding site for such a hydrophobic patch^35,36^. However, few Msp1 substrates contain similar hydrophobic patches, raising questions about the generality of a hydrophobic patch.

We propose that a hydrophobic mismatch between the substrate TMD and the lipid bilayer is the predominant mechanism for Msp1 substrate interactions. This model is particularly enticing as the predominant substrate for Msp1, mislocalized ER-TA proteins, are likely to exhibit a hydrophobic mismatch. Previous work has established that ER-TA proteins tend to be longer and more hydrophobic than mitochondrial TA proteins^37^. Therefore, mislocalized ER-TA proteins will likely have a hydrophobic mismatch with the OMM, leading to exposed hydrophobic residues that will drive Msp1 engagement^37^.

If the hydrophobic mismatch is the predominant means of substrate recognition, then Msp1 is likely engaging with the middle of the substrate rather than the terminus. This raises the question of how a membrane anchored substrate can gain access to the axial pore of the membrane anchored Msp1 ring. As previously proposed, there are two potential models that overcome this geometric problem^9,10^. First, a substrate could laterally diffuse into the axial pore through a gap in the Msp1 ring. Second, the Msp1 ring could assemble around a substrate. The ring assembly model is supported by biochemical data showing that WT Msp1 does not form a constitutive hexamer *in vitro* or *in vivo*^12,16^. The lateral diffusion model is supported by cryo-EM structures of ATPase inactive Msp1 hexamers in a lock-washer conformation with a seam that could support lateral diffusion of the substrate into the axial pore^35,36^.

A particularly attractive feature of the lateral diffusion model is that the cryo-EM structure of Msp1 shows an exposed hydrophobic groove in the N-domain of the Msp1 subunit the forms part of the seam in the lock washer^36^. Cross-linking studies have shown that this hydrophobic groove interacts with substrates^16^. As the N-domain is exposed to the lipid bilayer, this provides an obvious mechanism for simultaneously recognizing a hydrophobic mismatch and facilitating lateral diffusion of the substrate into the axial pore **(Figure 6G)**.

While our results demonstrate a prominent role for a hydrophobic mismatch in substrate recognition, they do not exclude the possibility of processive substrate unfolding driven by the Msp1 pore loops directly engaging with long, unstructured regions on the substrate terminus. Indeed, many AAA+ proteins have bipartite substrate recognition motifs, with long-unstructured regions showing non-sequence specific engagement with the pore loops^38,39^.

Our model also raises the possibility of Msp1 activity being regulated by changes in lipid composition of the OMM. Indeed, previous lipidomic work has shown that the OMM is one of the most fluid membranes in the cell, which should support robust Msp1 activity^40^. Furthermore, mitochondrial lipid composition exhibits daily oscillations^41^. Mitochondria also play a major role in lipid metabolism, with ER-Mitochondria contact sites facilitating the exchange of lipids between the two membranes^42^. Interestingly, Msp1 appears to preferentially engage with substrates at ER-Mitochondria contact sites^43^, suggesting that Msp1 may be particularly sensitive to changes in lipid metabolism. Future studies combining high resolution lipidomic work with *in vivo* Msp1 activity measurements^44^ will be important for enhancing our understanding of how changes in lipid composition regulate Msp1 activity.

In conclusion, we have developed a new assay to monitor the extraction of membrane proteins from a lipid bilayer by the AAA+ motor protein Msp1. Using this fully defined system, we systematically probed the role of the membrane composition and substrate TMD of Msp1 activity. We discovered that Msp1 recognizes substrates by a hydrophobic mismatch between the substrate TMD and the lipid bilayer. We also demonstrated that the rate limiting step in substrate extraction is removal of the TMD from the lipid bilayer. Together, this work provides foundational insights into how Msp1 and other membrane extractases perform the essential function of removing membrane proteins from a lipid bilayer.

## Materials and Methods

### Soluble Protein Purification *Δ1-32 Msp1*

Plasmids were transformed into *E. coli* BL21(DE3) containing a pRIL plasmid and expressed in terrific broth at 37°C until an OD600 of 0.6–1.0, cultures were induced with 0.25 mM IPTG and grown at room temperature for an additional 3–4 hr. Cells were harvested by centrifugation, and resuspended in Msp1 Lysis Buffer (20 mM Tris pH 7.5, 200 mM KAc, 20 mM imidazole, 0.01 mM EDTA, 1 mM DTT) supplemented with 0.05 mg/mL lysozyme (Sigma), 1 mM phenylmethanesulfonyl fluoride (PMSF) and 500 U of universal nuclease (Pierce), and lysed by sonication. The supernatant was isolated by centrifugation for 30 min at 4°C at 18,500 × *g* and purified by Ni-NTA affinity chromatography (Pierce) on a gravity column. Ni-NTA resin was washed with 10 column volumes (CV) of Msp1 Lysis Buffer and then 10 CV of Wash Buffer (Msp1 Lysis buffer with 30 mM imidazole) before elution with Lysis Buffer supplemented with 250 mM imidazole.

The protein was further purified by size exclusion chromatography (SEC) (Superdex 200 Increase 10/300 GL, GE Healthcare) in 20 mM Tris pH 7.5, 200 mM KAc, 1 mM DTT. Peak fractions were pooled, concentrated to 5–15 mg/mL in a 30 kDa MWCO Amicon Ultra centrifugal filter (Pierce) and aliquots were flash-frozen in liquid nitrogen and stored at –80°C. Protein concentrations were determined by A280 using a calculated extinction coefficient (Expasy).

#### SGTA and Calmodulin

GST-SGTA and GST-calmodulin were expressed as described above for soluble Msp1 constructs. Cells were harvested by centrifugation and resuspended in SGTA Lysis Buffer (50 mM HEPES pH 7.5, 150 mM NaCl, 0.01 mM EDTA, 1 mM DTT, 10% glycerol) supplemented with 0.05 mg/mL lysozyme (Sigma), 1 mM PMSF and 500 U of universal nuclease (Pierce), and lysed by sonication. The supernatant was isolated by centrifugation for 30 min at 4°C at 18,500 × *g* and purified by glutathione affinity chromatography (Thermo Fisher) on a gravity column. Resin was washed with 20 CV of SGTA Lysis Buffer and then eluted with 3 CV of SGTA Lysis Buffer supplemented with 10 mM reduced glutathione. The protein was further purified by SEC (Superdex 200 Increase 10/300 GL, GE Healthcare) in 20 mM Tris pH 7.5, 100 mM NaCl, 0.1 mM TCEP. Peak fractions were pooled, concentrated to 10 mg/mL in a 30 kDa MWCO Spin Concentrator (Pierce) and aliquots were flash-frozen in liquid nitrogen and stored at –80°C. Protein concentrations were determined by A280 using a calculated extinction coefficient (Expasy).

#### LgBiT and MBP-Ubiquitin

LgBiT and MBP-Ubiquitin were expressed as described above for soluble Msp1 constructs. Cells were harvested by centrifugation and resuspended in Lysis Buffer (20 Tris 7.5, 200 NaCl, 1 DTT, 0.01 EDTA, 20 mM imidazole) supplemented with 0.05 mg/mL lysozyme (Sigma), 1 mM PMSF and 500 U of universal nuclease (Pierce), and lysed by sonication. The supernatant was isolated by centrifugation for 30 min at 4°C at 18,500 × *g* and purified by Ni-NTA affinity chromatography (Pierce) on a gravity column. Ni-NTA resin was washed with 10 column volumes (CV) of Msp1 Lysis Buffer and then 10 CV of Wash Buffer (20 Tris 7.5, 200 NaCl, 1 DTT, 0.01 EDTA, 30 mM imidazole) before elution with Lysis Buffer supplemented with 250 mM imidazole. The protein was further purified by size exclusion chromatography (SEC) (Superdex 200 Increase 10/300 GL, GE Healthcare) in 20 mM Tris pH 7.5, 200 mM NaCl, 1 mM DTT. Peak fractions were pooled, concentrated to 5–15 mg/mL in a 30 kDa MWCO Amicon Ultra centrifugal filter (Pierce) and aliquots were flash-frozen in liquid nitrogen and stored at –80°C. Protein concentrations were determined by A280 using a calculated extinction coefficient (Expasy).

### Membrane Protein Purification

#### SumoTMD substrates

The main substrate (SUMO-Sec22 TMD) is in a pET28a plasmid backbone. Cloning of all synthetic TMD substrates was conducted via traditional PCR/restriction enzyme ‘cut and paste’ methods to swap out Sec22 TMD and verified by Sanger sequencing. The layout of all constructs is His_6_-3C protease site-3xFlag-SUMO-*TMD*-thrombin protease site-HiBiT tag. The DHFR-Sec22 substrate has the same set up except the SUMO domain is replaced with mouse DHFR. Expression and purification of the DHFR-Sec22 substrate was the same as the Sumo-Sec22 substrate.

After transforming the SumoTMD plasmid into *E. coli* BL21(DE3)/pRIL, the cells were grown in terrific broth at 37°C until an OD_600_ of 0.6-0.8, and then induced with 0.25 mM IPTG and grown at room temperature for an additional 3-4 hr. Cells were harvested by centrifugation, and resuspended in SumoTMD Lysis Buffer (50 mM Tris pH 7.5, 300 mM NaCl, 10 mM MgCl_2_, 10 mM Imidazole, 10% glycerol) supplemented with 0.05 mg/mL lysozyme (Sigma), 1 mM PMSF and 500 U of benzonase (Sigma), and lysed by sonication. Membrane proteins were solubilized by addition of n-dodecyl-β-D-maltoside (DDM) to a final concentration of 1% and rocked at 4°C for 30’. Lysate was cleared by centrifugation for at 4°C for 1 hr at 35,000 x g and purified by Ni-NTA affinity chromatography.

Ni-NTA resin was washed with 10 column volumes (CV) of SumoTMD Wash Buffer 1 (50 mM Tris pH 7.5, 500 mM NaCl, 10 mM MgCl_2_, 10 mM imidazole, 5 mM β-mercaptoethanol (BME), 10% glycerol, 0.1% DDM). Resin was then washed with 10 CV of SumoTMD Wash Buffer 2 (same as Wash Buffer 1 except with 300 mM NaCl and 25 mM imidazole) and 10 CV of SumoTMD Wash Buffer 3 (same as Wash Buffer 1 with 150 mM NaCl and 50 mM imidazole) and then eluted with 3 CV of SumoTMD Elution Buffer (same as Wash Buffer 3 except with 250 mM imidazole).

The protein was further purified by size exclusion chromatography (SEC) (Superdex 200 Increase 10/300 GL, GE Healthcare) in 50 mM Tris pH 7.5, 150 mM NaCl, 10 mM MgCl_2_, 5 mM BME, 10% glycerol, 0.1% DDM. Peak fractions were pooled, aliquots were flash-frozen in liquid nitrogen and stored at –80°C. Protein concentrations were determined by A_280_ using a calculated extinction coefficient (Expasy).

### Liposome preparation

Liposomes mimicking the lipid composition of the yeast OMM, as well as the thickness and fluidity series, were prepared as described previously^7^. Briefly, a lipid film was prepared by mixing chloroform stocks of lipids applicable to each type of liposome:

**Nickel liposomes:** chicken egg phosphatidyl choline (Avanti 840051C), chicken egg phosphatidyl ethanolamine (Avanti 840021C), bovine liver phosphatidyl inositol (Avanti 840042C), synthetic DOPS (Avanti 840035C), synthetic TOCL (Avanti 710335C), and 1,2-dioleoyl-*sn*-glycero-3-[*N*-((5-amino-1-carboxypentyl)iminodiacetic acid)succinyl] Nickel salt (Avanti 790404) at a 48:28:10:8:4:2 molar ratio with 1 mg of DTT.

**T1:** synthetic 14:1, 14:1 phosphatidyl choline (Avanti 850346C) and synthetic 18:1, 18:1 1,2-dioleoyl-*sn*-glycero-3-[*N*-((5-amino-1-carboxypentyl)iminodiacetic acid)succinyl] Nickel salt (Avanti 790404) at a 98:2 molar ratio with 1 mg of DTT.

**T2:** synthetic 16:1, 16:1 phosphatidyl choline (Avanti 850358C) and synthetic 18:1, 18:1 1,2-dioleoyl-*sn*-glycero-3-[*N*-((5-amino-1-carboxypentyl)iminodiacetic acid)succinyl] Nickel salt (Avanti 790404) at a 98:2 molar ratio with 1 mg of DTT.

**T3:** synthetic 18:1, 18:1 phosphatidyl choline (Avanti 850375C) and synthetic 18:1, 18:1 1,2-dioleoyl-*sn*-glycero-3-[*N*-((5-amino-1-carboxypentyl)iminodiacetic acid)succinyl] Nickel salt (Avanti 790404) at a 98:2 molar ratio with 1 mg of DTT.

**T4:** synthetic 20:1, 20:1 phosphatidyl choline (Avanti 850396C) and synthetic 18:1, 18:1 1,2-dioleoyl-*sn*-glycero-3-[*N*-((5-amino-1-carboxypentyl)iminodiacetic acid)succinyl] Nickel salt (Avanti 790404) at a 98:2 molar ratio with 1 mg of DTT.

**T5:** synthetic 22:1, 22:1 phosphatidyl choline (Avanti 850398C) and synthetic 18:1, 18:1 1,2-dioleoyl-*sn*-glycero-3-[*N*-((5-amino-1-carboxypentyl)iminodiacetic acid)succinyl] Nickel salt (Avanti 790404) at a 98:2 molar ratio with 1 mg of DTT.

**F1:** synthetic 18:1, 18:1, *trans* phosphatidyl choline (Avanti 850376C) and synthetic 18:1, 18:1 1,2-dioleoyl-*sn*-glycero-3-[*N*-((5-amino-1-carboxypentyl)iminodiacetic acid)succinyl] Nickel salt (Avanti 790404) at a 98:2 molar ratio with 1 mg of DTT.

**F2:** synthetic 18:0, 18:1 phosphatidyl choline (Avanti 850467C) and synthetic 18:1, 18:1 1,2-dioleoyl-*sn*-glycero-3-[*N*-((5-amino-1-carboxypentyl)iminodiacetic acid)succinyl] Nickel salt (Avanti 790404) at a 98:2 molar ratio with 1 mg of DTT.

**F3:** synthetic 18:1, 18:1, *cis* phosphatidyl choline (Avanti 850375C) and synthetic 18:1, 18:1 1,2-dioleoyl-*sn*-glycero-3-[*N*-((5-amino-1-carboxypentyl)iminodiacetic acid)succinyl] Nickel salt (Avanti 790404) at a 98:2 molar ratio with 1 mg of DTT.

**F4:** synthetic 18:2, 18:2 phosphatidyl choline (Avanti 850385C) and synthetic 18:1, 18:1 1,2-dioleoyl-*sn*-glycero-3-[*N*-((5-amino-1-carboxypentyl)iminodiacetic acid)succinyl] Nickel salt (Avanti 790404) at a 98:2 molar ratio with 1 mg of DTT.

Chloroform was evaporated under a gentle steam of nitrogen and then left on a vacuum (<1 mTorr) overnight. Lipid film was fully resuspended in Liposome Buffer (50 mM HEPES KOH pH 7.5, 15% glycerol, 1 mM DTT) to a final concentration of 20 mg/mL and then subjected to five freeze-thaw cycles with liquid nitrogen. Liposomes were extruded 15 times through a 100 nm filter at 60°C, distributed into single-use aliquots, and flash-frozen in liquid nitrogen.

### Reconstitution into Liposomes

For extraction assays proteoliposomes were prepared by mixing 2 μM TA protein (SumoTMD), and 2 mg/mL of Nickel liposomes in Reconstitution Buffer. Detergent was removed by adding 25 mg of biobeads and rotating the samples for 16 hr at 4°C. Material was removed from biobeads and added to fresh 25 mg biobeads and rotated for another 1-3 hr at RT. Unincorporated TA protein was pre-cleared by incubating the reconstituted material with excess (5 μM) GST-SGTA and GST-calmodulin and passing over a glutathione spin column (Pierce #16103); the flow through was collected and used immediately for dislocation assays.

### C-Laurdan Spectroscopy

C-laurdan spectroscopy was performed by using 10 μL of pre-formed F-series liposomes and 90 μl of extraction buffer (50 mM Hepes pH 7.5, 200 mM KAc, 7 mM MgAc, 2 mM DTT, 10 μm Ca^2+^) to match overnight reconstitution volume. Final 50 μg/ml of C-laurdan dye (Fisher) was added and a fluorescence spectrum was read. The sample was excited at 375 nm and an emission spectrum from 400 to 600 nm with 3 nm step size was recorded. The GP value (-1 most disordered, +1 most ordered membranes) was measured as described previously^45^ by integrating the intensities between 400 and 460 nm (*I*_Ch1_), and 470 and 530 nm (*I*_Ch2_): GP = (*I*_CH1_*-I*_CH2_) / (*I*_CH1_*+ I*_CH2_).

### Extraction Assay

Extraction assays contained 30 μL of pre-cleared proteoliposomes, 5 μM GST-SGTA, 5 μM GST-calmodulin, 3 μM Msp1, 8 μM MBP-Ubiquitin, 1 mg/mL bovine serum albumin (sigma), and 110 mM ATP and the final volume was adjusted to 100 μL with Extraction Buffer (50 mM HEPES KOH pH 7.5, 200 mM potassium acetate, 7 mM magnesium acetate, 2 mM DTT, 0.1 μM calcium chloride). Samples were incubated at 30°C for 30 minutes and then transferred to ice. Samples were transferred to polycarbonate centrifuge tubes (Beckman-Coulter #343776) and centrifuged at 150,000 x *g* in TLA-120.1 Fixed-Angle Rotor (Beckman-Coulter) for 30’ at 4°C. The top 20 ul was collected and combined with 1.5 μM LgBiT and the final volume was adjusted to 50 μL with Extraction Buffer. After incubating at RT for 25 minutes, samples were plated onto white 96-well plates (Corning #07201204) and 20 μL of lytic furimazine (Promega #N3030) was added. Immediately plates were read at 480 nm wavelength to measure luminescence.

Extraction assays with DHFR-Sec22 were performed as described above except DMSO or methotrexate were added 5 minutes prior to initiation of the assay with ATP. A 1 mM methotrexate stock solution was prepared in DMSO and then added to at a final concentration of 5 μM. The final concentration of DMSO in the assay was 0.5% of the reaction volume.

### Standard curve measurements

Standards were made starting with 20% of the volume of pre-cleared liposomes used in the extraction and brought up to 100 μL with Extraction Buffer (50 mM HEPES KOH pH 7.5, 200 mM potassium acetate, 7 mM magnesium acetate, 2 mM DTT, 0.1 μM calcium chloride). This was then diluted to make 15%, 10%, and 5% samples. 20 μL from each was then combined with 1.5 μM LgBiT and the final volume was adjusted to 50 μL with Extraction Buffer. After incubating at RT for 25 minutes, samples were plated onto white 96-well plates (Corning #07201204) and 20 μL of lytic furimazine (Promega #N3030) was added. Immediately plates were read at 480 nm wavelength to measure luminescence.

### Protease Protection Assay

Protease protection assay was carried out in a 10 μL total volume with 7 μL of pre-cleared liposomes, 2 U of thrombin protease, and 1% of Triton X-100 (where indicated). Samples were incubated at RT for 1 hour and then 1 μL of 200 mM PMSF was added. The samples were reverse quenched into 90 μL of boiling 1% SDS and incubated at 95°C for 10 minutes.

## Acknowledgements

The authors wish to thank members of the Wohlever and Tristam-Nagel labs for helpful discussions and feedback on the project.

## Author Contributions

Conceptualization: HLF, DG, MLW; Methodology: HLF, DG, BS, BA, MLW; Investigation: HLF, DG, BS, BA; Writing – Original Draft: HLF, MLW; Writing – Review and Editing: HLF, DG, BS, BA, MLW.

